# Systems-level longitudinal immune profiling reveals individualized immunotypes and genetic associations

**DOI:** 10.64898/2026.03.21.713378

**Authors:** Alberto Zenere, Xingyue Wang, Ziyang Tan, Tadepally Lakshmikanth, Jaromir Mikes, Yang Chen, Maria Johansson, Göran Bergström, Fredrik Edfors, Mathias Uhlén, Petter Brodin, Wen Zhong

## Abstract

The human immune system exhibits substantial inter-individual variation, yet how this variability is coordinated across the system and persists over time remains incompletely defined. Here, we integrated whole-genome sequencing with longitudinal multi-omics immune profiling in 101 healthy individuals followed over two years, combining mass cytometry-based immune cell profiling, PBMC transcriptomics, plasma proteomics, and clinical measurements. Coordinated variation across cellular composition, gene expression, and circulating proteins revealed reproducible immune patterns within individuals across visits. Integrative analyses identified three major immunotypes: adaptive lymphoid (CD4⁺ T cell and B cell enriched), myeloid-inflammation (monocyte and dendritic cell-driven), and cytotoxic (CD8⁺ T cell-dominated), each associated with distinct metabolic and inflammatory profiles. Genome-wide association analyses identified cell-type-specific quantitative trait loci primarily affecting memory lymphocytes, and a polygenic score for memory B cells correlated with both cellular abundance and transcriptional activity. Together, these findings provide a systems-level view for understanding baseline immune heterogeneity in human populations.

**Teaser:** Longitudinally stable immune individuality reflects coordinated modules shaped by systemic physiology and genetics

## Introduction

The human immune system exhibits significant variability across individuals. Each individual maintains a distinct immunological profile, or immunotype, characterized by the composition and activity of immune cells ^1–3^, as well as soluble mediators such as cytokines, chemokines, antibodies, and complement proteins. These individualized immunotypes are increasingly recognized as key determinants of susceptibility to infections ^4,5^, responsiveness to vaccination ^6–11^, and the efficacy of immune-targeted therapies ^12–14^. Understanding the variability and underlying mechanisms of individual immunotypes, and how they respond to physiological and environmental perturbations, is therefore essential to precision immunology, which aims to tailor prevention and treatment strategies to each individual’s immune landscape ^15^.

The inter-individual immune variability is shaped by a complex interplay of genetic, physiological, and environmental factors. Advances in high-throughput technologies, such as high-throughput sequencing, mass cytometry (CyTOF), single-cell omics, and multiplex proteomics, have significantly enhanced our ability to profile individual immunotypes at the systems level. These approaches have revealed that the range of variation in major immune cell populations such as B cells, CD4⁺ T cells, CD8⁺ T cells, and NK cells can vary by many orders of magnitude among healthy individuals ^1^. Recent longitudinal studies further showed that several cytokines and chemokines, including IL-6, IFN, IL-8, also vary widely across individuals, supporting the concept of a personal immune “set point” maintained through coordinated regulation ^16–18^.

Given the complexity of the human immune system, the relative contributions of different factors vary significantly across immune parameters. Genome-wide associations have revealed significant genetic influences on immune system variability. For instance, heritability estimates for key immune traits, particularly within the adaptive immune system, range from 3% to 87%, with T cell and B cell populations showing substantial heritability ^19–21^. In addition, numerous important genetic variants have been identified to influence HLA, cytokine levels, and immune cell gene expressions ^22,23^, emphasizing the substantial role of genetic regulation in immune variability. Conversely, a twin study showed that most immune parameters (> 70% of the immune traits) are dominated by non-heritable influences ^18^. Numerous studies have shown that the immune variability is also shaped by factors such as age, sex and cohabitation ^1,24^.

Despite these advances, substantial gaps remain in understanding how immune individuality is organized across molecular and cellular layers, and to what extent it is influenced by heritable and non-heritable factors. The immune system functions as an integrated network in which genetic architecture, cellular interactions, and environmental exposures collectively shape each person’s unique immunological state. Understanding how these layers collectively define an individual’s system-level immunotype, and how genetic architecture and non-genetic exposures interact to jointly shape and stabilize these personalized immune states, is crucial for monitoring the dynamic changes in immune system components in response to external influences and internal physiological changes, providing a more comprehensive understanding of individual immune variability.

In this study, we present a systems-level view of human immune individuality with the longitudinal integrative analysis of whole-genome sequencing with multi-omics immune profiling in 101 individuals over two years. Distinct and stable immunotypes were identified, characterized by coordinated cellular, transcriptional, and proteomic signatures. These immunotypes were further associated with genetic variations and systemic clinical and biochemical markers, providing an integrated understanding of how inherited and physiological factors jointly shape the diversity and stability of human immune individuality.

## Results

### The Swedish SciLifeLab SCAPIS Wellness Profiling cohort

In this study, we examined 101 clinically healthy individuals enrolled in the Swedish SciLifeLab SCAPIS Wellness Profiling (S3WP) program. Participants were followed longitudinally for two years with repeated assessments of their molecular and clinical profiles ^1,25–27^, see **Figure 1a, b**. The participants were evenly distributed by sexes (53 females, 51%) and were between 50 to 66 years of age (**Figure 1c, d**). At each visit, extensive anthropometric and biochemical measurements were performed, with a total of 29 clinical variables were included in the analysis, see **Supplementary Table 1**. Peripheral blood mononuclear cells (PBMCs) and plasma samples were obtained at each visit for multi-omics profiling. PBMC immune cell profiling was analyzed by CyTOF, resolving 53 immune cell subsets across 18 major cell types, including NK cells, monocytes and dendritic cells, basophils, innate lymphoid cells (ILCs), and naïve and memory B cells and T cells (**Figure 1e, Supplementary Table 1**). Transcriptomics of PBMCs were analyzed and obtained 16,985–18,100 expressed genes (TPM > 0) across 528 PBMC samples (detailed in Methods). To assess the quality of the PBMC transcriptomics data, we compared the PBMC transcriptomic profiles from the S3WP study with the PBMC and 18 flow-sorted immune cell transcriptomes from the Human Protein Atlas (HPA) ^28^; as expected, the PBMC samples from the S3WP project were clustered together with PBMC samples from the HPA (**Figure 1g**). Plasma proteome profiling was performed using Olink proximity extension assays, quantifying 794 plasma proteins covering immune, inflammatory, cardiovascular, metabolic, and organ damage related biological processes. In addition, whole genome sequencing was performed for all participants in the S3WP study, identifying approximately 6.7 million high-confidence single nucleotide polymorphisms (SNPs) per individual, enabling the analyses of genetic influence on immune variation.

**Figure 1:**
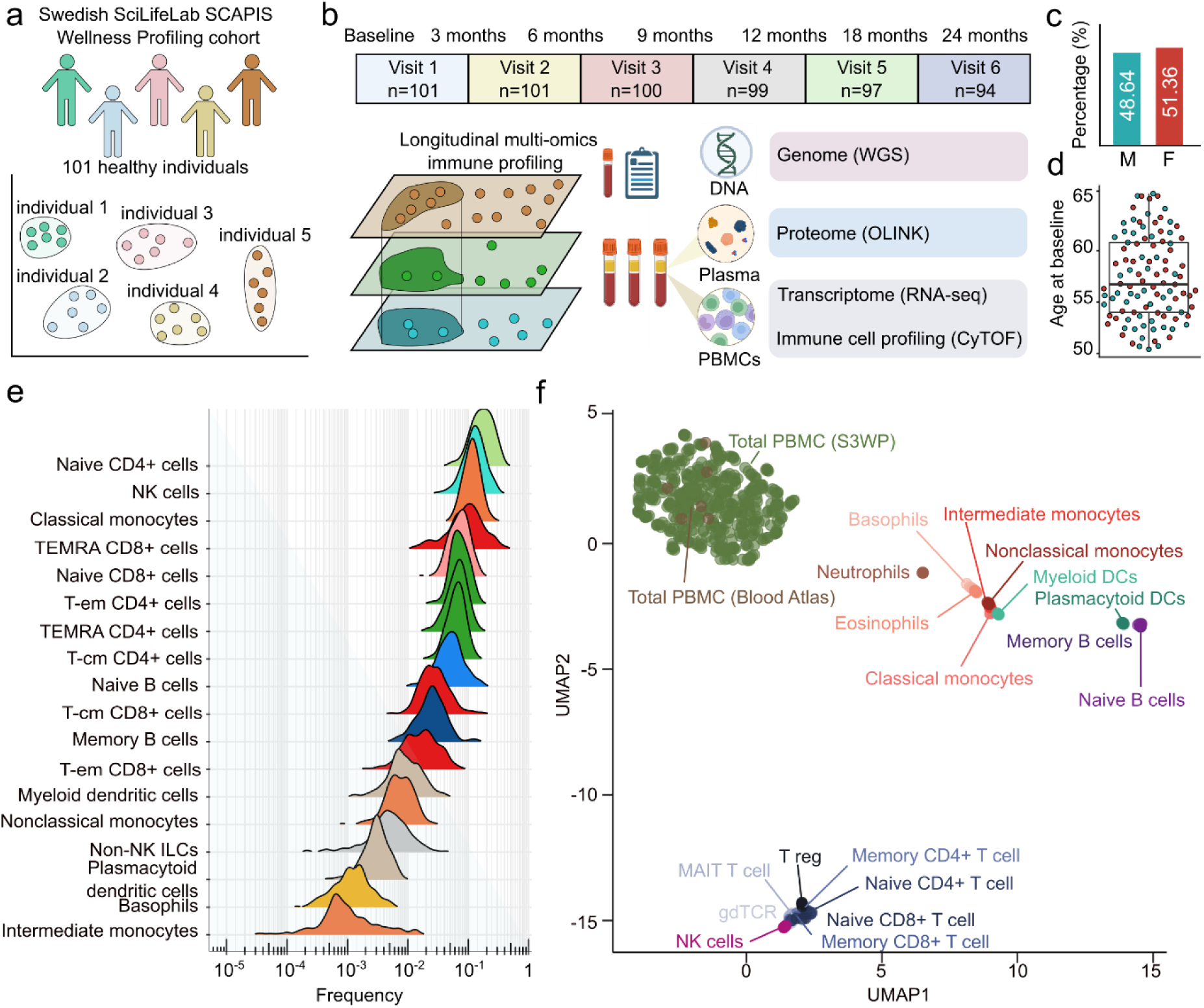
Overview of study design. (a) The S3WP cohort comprises 101 clinically healthy individuals characterized by longitudinal multi-omics profiling. Repeated measurements from the same individual enabled assessments of intra-and inter-individual immune variation and cross-omic integration. (b) The participants were followed longitudinally across six visits over two years. At each visit, PBMCs and plasma samples were collected for CyTOF, RNA-seq, and Olink proteome profiling; WGS was performed at baseline. (c) Sex distribution of the participants. (d) Age distribution at enrollment. (e) Frequency distributions of 18 major immune cell populations derived from CyTOF analysis across all samples. (f) UMAP clustering of PBMC transcriptomics profiles from the S3WP cohort alongside immune-cell transcriptomics from the Human Protein Atlas. Colors indicate immune cell populations.

### Stable and individualized systemic immune profiles

To evaluate systemic immune individuality in the S3WP cohort, we compared the sample similarity across three omics layers, including the immune cell profiling, PBMC transcriptomics, and plasma proteome profiling using t-distributed stochastic neighbor embedding (t-SNE) see **Figure 2a-d**. All three layers exhibited substantial inter-individual variability; however, the degree of individual segregation differed across modalities. PBMC transcriptomics and plasma proteomics revealed strong individual-level segregation (**Figure 2b and 2d**), whereas immune cell profiling showed more moderate individual-specific clustering (**Figure 2a**). These findings indicated that molecular immune features captured highly personalized signatures that exceeded the variation observed at the cellular level. Hierarchical clustering of PBMC transcriptomes from S3WP participants and reference immune cell types from the Human Protein Atlas further confirmed this immune individuality, with PBMC samples forming distinct individual-specific clusters and clearly separating from other immune lineages (**Figure 2e**). To assess the long-term stability of individual PBMC transcriptomics, we integrated 174 additional PBMC transcriptomics data from the same individuals in the S3WP project that were collected in the second year after the initial visit (visits 5 and 6). Euclidean distance analyses of PBMC transcriptomics within and between individuals showed that intra-individual distances remained consistently and significantly lower than inter-individual distances (Wilcoxon test, *P* < 2.2 × 10⁻¹⁶), with no detectable difference between same-year and cross-year comparisons (**Figure 2f**). Moreover, each individual’s transcriptomics profile at visit 6 was closest to their own first-year samples, indicating that individual immune signatures were stable over time (**Figure 2g**).

**Figure 2.**
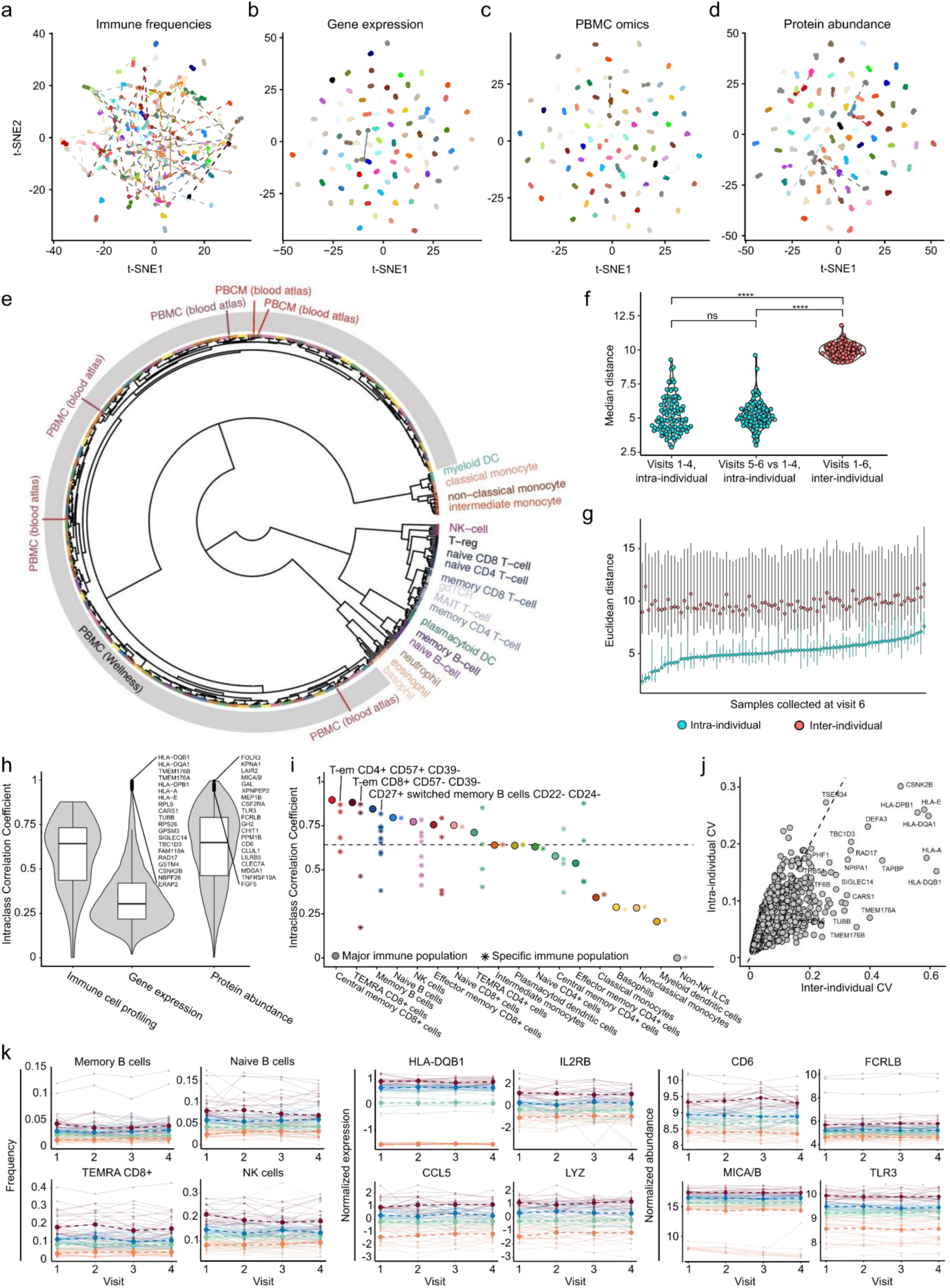
Individual immune signatures. (a-d) t-SNE performed on (a) PBMC immune frequencies, (b) PBMC gene expression, (c) immune frequencies and gene expression combined, and (d) plasma proteomics. (e) Clustering of transcriptomic data from our study and the Human Protein Atlas. (f) Distribution of Euclidean distance in transcriptomic profiles between collected samples (i) from the same participant within the first year (visit 1-4), (ii) from the same participant between year 1 and year 2 (visit 5-6), and (iii) from different participants. Wilcoxon tests were used for statistical analysis (ns, not significant; ****, *P* < 0.0001). (g) Euclidean distance between each transcriptomic profile from the last visit and the sample collected during the first year, divided into (i) pairs of samples from the same individual (light blue) and (ii) pairs of samples from different participants (red). (h) Distribution of intraclass correlation coefficient of immune cell profiling, transcriptomic and plasma proteomics. (i) Intraclass correlation coefficient of all 53 immune populations. (j) Intra- and inter-individual coefficients of variation of gene expression. (k) Examples of genes, immune populations and proteins showing individual expression profiles. Samples are colored according to their expression levels at visit 1 (orange: 0-25%, green: 25–50%, blue: 50–75%, purple: 75–100%). The highlighted dots indicate the median level of the corresponding group at that visit.

### Key immune features driving inter-individual diversity

To identify immune features contributing most to inter-individual variability, we calculated the Intraclass Correlation Coefficient (ICC) for all immune cell types, expressed genes in PBMCs, and plasma proteins using linear mixed-effects models (LME) with sex and age as fixed effects, **Supplementary Table 2**. Although on sample-level transcriptomic and proteomic profiles exhibited stronger individual segregation in low-dimensional embedding analyses, feature-level stability was greater for immune cell frequencies and plasma proteins than for PBMC gene expression (**Figure 2h)**. Specifically, adaptive immune cells showed higher inter-individual variation than innate immune cells (**Figure 2i**). Among them, memory CD4+ subsets (T-em CD57+ CD39-, TEMRA CD57+ CD39-), memory CD8+ (T-cm CD57-CD39-, T-cm CD57+ CD39-, TEMRA CD57-CD39-, TEMRA CD57+ CD39-) and memory B cell types (CD27+ switched memory B cells CD22-CD24-) showed the highest ICC values (ICC=0.80-0.88), **Figure 2i**.

At the transcriptional level, 2,544 genes exhibited greater inter-individual variation than intra-individual variability (**Figure 2j**). Among them, 39 genes showed exceptionally high inter-individual variance (ICC > 0.90, n = 39). These included genes involved in antigen repertoire shaping (*ERAP2, TAPBP, PSMB9*), antigen presentation (*HLA-A*, *HLA-B*, *HLA-E*, *HLA-DPB1*, *HLA-DQA1*, *HLA-DQB1*), cytotoxic effector function (*CD8A, GZMH, GNLY)* and modulators of the NLRP3 inflammasome (*TMEM176A*, *TMEM176B*, *SIGLEC14*), see **Figure 2k** and **Supplementary Table 2**. Pathway enrichment analysis of genes with high inter-individual variation highlighted immune processes linked to mononuclear cell activation and proliferation, (**Supplementary Figure 1a**), while gene set enrichment analysis (GSEA) based on ICC ranking showed enrichment in humoral responses, myeloid activation, and cytotoxic pathways, **Supplementary Figure 1b**.

To systematically assess variability across immune pathways, we examined ICC levels of genes within 26 curated immune gene sets (**Supplementary Table 2**). Interestingly, a clear pattern was observed that receptor genes, including HLA molecules, the T cell receptors (TCRs), B cell receptors (BCRs), natural killer group 2 (NKG2), and C-type lectin receptors (CLRs), showed the greatest inter-individual variability, with ICC values consistently above the genome-wide median, **Supplementary Figure 1c**. Moderate inter-individual variability was observed in genes encoding secreted effector molecules, such as cytotoxic mediators (*GZMA*, *GZMB*, *GZMH*, *GZMK*, *GZMM*, *PRF1*) and antimicrobial proteins (*LYZ*, *S100A8*, *S100A9*), **Supplementary Figure 1c**. In contrast, intracellular signaling components, including T cell signaling genes, interferon regulatory factors (IRFs) and inflammasome-related genes, exhibited ICC values close to the genome-wide median level, **Supplementary Figure 1c**. Overall, receptor-encoding genes exhibited the strongest individual specificity, followed by genes encoding secreted and intracellular proteins **Supplementary Figure 1d**.

We next examined the contribution of sex and age to transcriptional variation. The results revealed that sex was associated with differential expression of 23% of genes (n = 1,950; FDR < 0.05), of which 1,264 were upregulated in females and 686 in males (**Supplementary Figure 1e)**. Among the genes upregulated in females, the strongest sex-bias was found in females with known X-chromosome inactivation (XCI) escape genes, such as *KDM5C, KDM6A, CXorf38, DDX3X, RPS4X,* and *EIF1AX*. Notably, 27 sex-biased genes also showed high inter-individual variability (ICC > 0.8). Among these, one notable example was *IL4R,* which was upregulated in females, in line with previous reports that females tended to exhibit stronger IL4 signaling ^29^ (**Supplementary Figure 1f).** At the pathway level, sex-biased expression patterns were also evident, with higher expression of NKG2 receptors, AIM2-like receptors, and Toll-like receptors in males, and increased expression of B cell receptor–related genes in females (**Supplementary Figure 1g**). In contrast, no significant associations between gene expression and age were detected after multiple testing correction, possibly due to the relatively narrow age range of the cohort (50–65 years).

At the plasma protein level, we identified an additional layer of immune individuality captured through circulating mediators, including innate pattern recognition and activation markers (TLR3, CLEC7A), stress and cytotoxicity-associated ligands (MICA/B), and modulators of myeloid–lymphoid communication (CSF2RA). These circulating proteins provided complementary evidence that systemic immune signaling, beyond cellular composition and gene expression, was also personalized across individuals, particularly in antigen processing and cytotoxic pathways, **Figure 2j**. Significant correlations (Spearman’s ρ > 0.2) between plasma proteomics and CyTOF profiling revealed a close link between immune cell composition and the circulating milieu, highlighting proteins associated with the activation of specific cell types. Notable associations included TCL1A, FCER2, FCRL1, CD28, APBB1IP, LGALS1 and SELL with naïve B cells; GZMA, GZMH, FCRL5/6, KLRD1, IL7R, SELPLG, SLAMF1, and ADGRG1 in CD8+ TEMRA T cells; TREM1, IL1RL1, IL6 with classical monocytes and myeloid DCs; and GZMB, NQO2, and CHRDL2 with NK cells (**Supplementary Table 2**).

### Integrative analysis of individualized immune cell and gene expression network

To investigate genome-wide associations between immune cell populations and PBMC gene expressions, we applied linear regression models relating isometric log ratio (ILR) transformed cell frequencies to gene expressions ^30^, with age and sex included as covariates and between-visit variability modeled as a random effect (see Methods for details). In total, 2,966 significant cell-gene associations were identified (FDR = 0.05); most of which were linked to classical monocytes (n = 988), TEMRA CD8+ T cells (n = 425), NK cells (n = 337), and memory B cells (n = 247), see **Figure 3a** and **Supplementary Table 3**. Notable associations included *BANK1*, *BLK* and *CD79A* with memory B-cells; *CD163*, *CD14* and *CLEC4E* with classical monocytes; and *GLNY*, *GZMB* and *KLRD1* associated with NK cells, **Figure 3b**. These genes represented canonical markers of their respective immune populations and had previously been shown to exhibit cell type-specific expression in the Human Protein Atlas ^28^ (**Supplementary Figure 2a-c**). Several of the identified associations between immune cell-type and PBMC gene expressions could be validated by benchmarking against known cell type markers retrieved from Azimuth ^31^ and DICE (Database of Immune Cell Expression, eQTLs, and Epigenomics) ^32^, see **Supplementary Figure 2d-f**. Further, the associations were validated by testing the overlap with the Human Protein Atlas ^28^, see **Figure 3c**.

**Figure 3.**
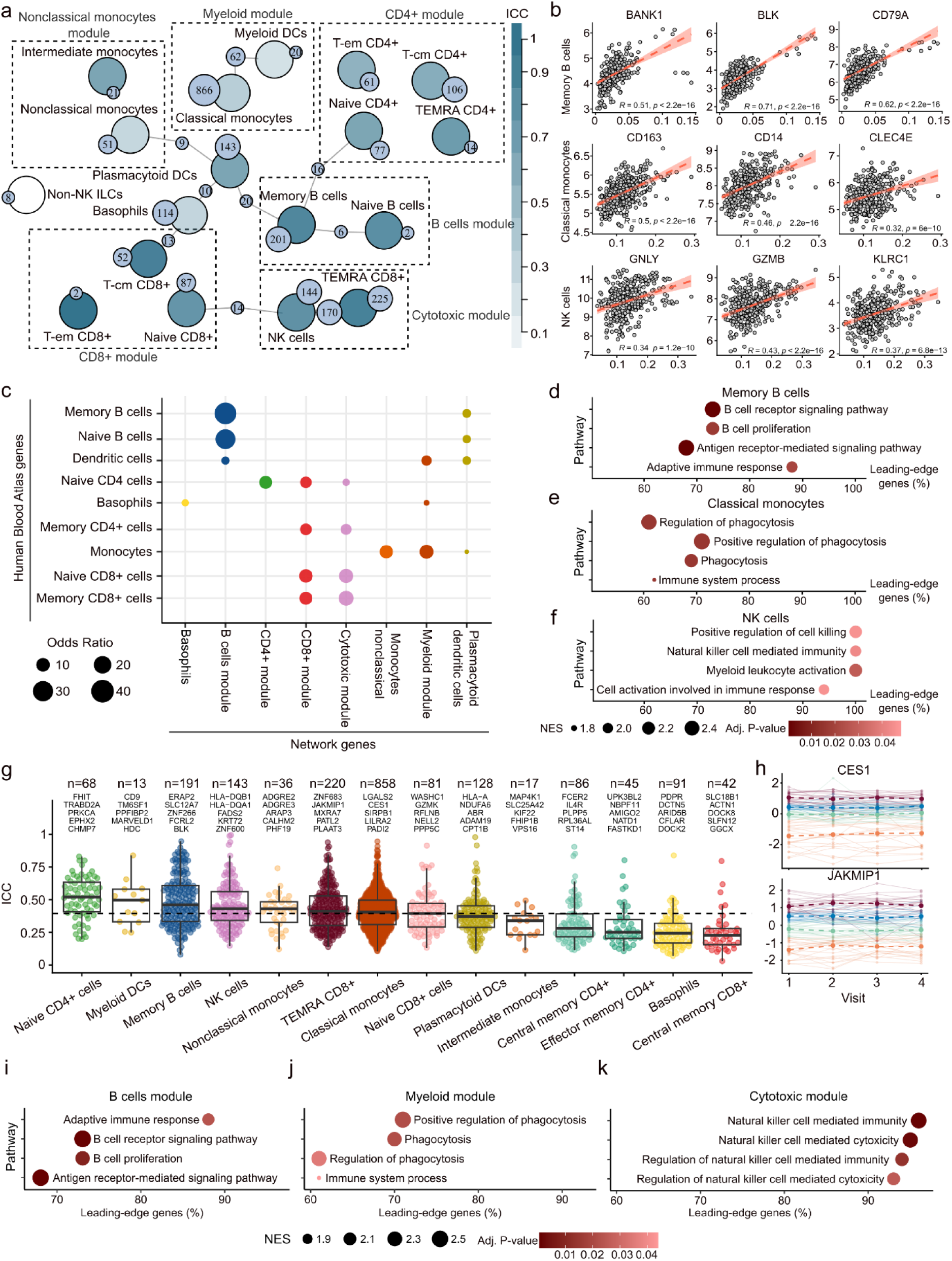
PBMC association network. (a) Summary of significant associations (FDR = 0.05) between immune frequencies and gene expression. The cell types are organized into six distinct modules, associated with similar immune functions. Each cell type is colored based on the ICC value of its frequency. (b) Examples of significant associations between immune cell composition and gene expression; Pearson correlation coefficients are indicated. (c) Statistically significant overlap (Fisher’s exact test, adjusted using Benjamini-Hochberg procedure) between the PBMC network associations and the annotation from the Human Protein Atlas. (d-f) Top four significant GSEA results, for memory B cells, classical monocytes and NK cells, respectively. (g) Distribution of ICCs across 18 major immune populations; genes with the top 5 highest ICC are labeled. On top is the number of genes uniquely associated with each cell type. (h) Two examples of genes with high ICC associated with immune cell types. Samples are colored according to their expression levels at visit 1 (orange: 0–25%, green: 25–50%, blue: 50–75%, purple: 75–100%) (i-k) Top four significant GSEA results the cytotoxic, B cells and myeloid module, where genes were ordered according to their ICC. GSEA, gene set enrichment analysis, central memory, T-cm; effector memory, T-em; dendritic cells, DCs; intraclass correlation coefficient, ICC.

To examine the individual variability and relevant gene functions within each cell type, GSEA was performed using ICC to rank genes by their between-individual variation. The results highlighted that genes with higher ICC tended to be involved in cell type-specific functions, **Figure 3d-f**. Consistently, we observed that many high-ICC genes were also cell-type specific, such as *CES1* with classical monocytes, and *JAKMIP1* with TEMRA CD8+ T cells, **Figure 3g and 3h**. Several of these high ICC genes were involved in core cell type specific functions, such as B-cell receptor signaling and identity in B cells (*BLK, FCRL2*), activation (*ADGRE2/*3, *LILRA2*), phagocytosis (SIRPB1), lipid metabolism (*CES1*) in monocytes and terminal effector differentiation in TEMRA CD8+ T cells (*ZNF683*).

In addition to cell-type specific gene associations, we observed connections across different cell-types. For example, TEMRA CD8+ T cells and NK cells shared a substantial set of associated genes (n = 170, Fisher’s exact test *P* < 2.2e-16, odds ratio = 31), which were enriched for cytotoxic pathways (**Supplementary Figure 2g**). Based on the immune cell-gene network analysis, we grouped PBMC cell populations into nine modules, including nonclassical/intermediate monocyte module, classical monocyte/mDC module, CD4+ T cell module, CD8+ T cell module, cytotoxic module, B cell module, plasmacytoid DCs (pDCs) module, basophil module and innate lymphoid cell module (**Figure 3a**). These modules were consistent with the clustering patterns derived from CyTOF surface-marker expressions in the S3WP cohort (see **Supplementary Figure 2h, i**). Among the nine modules, the CD4+ T cell, CD8+ T cell, B cell, and cytotoxic modules showed highest inter-individual variability, as shown by GSEA (using ICC as metric to order the genes) on genes involved in each module, which pointed to pathways related to module-specific cell types, see **Figure 3i-k** and **Supplementary Table 3**.

### Clustering analysis revealed three major distinct and stable immunotypes

To investigate the immune heterogeneity across individuals, we performed sample-level clustering of the immune cell-gene network using K-nearest neighbor (KNN), and Louvain algorithm. Although the samples exhibited a continuum of immune variation as reported previously ^3,33^, this analysis revealed three distinct immunotypes (clusters), see **Figure 4a**. These clusters showed significant differences in immune cell composition (**Figure 4b**, **c** and **Supplementary Figure 3a**). Cluster A consisted predominantly of female samples (n = 73, 73%) and was characterized by higher frequencies of B cells and CD4+:CD8+ ratio. Cluster B comprised mainly of male samples (n = 137, 70%) and showed increased frequencies of innate immune cell populations, including monocytes and dendritic cells. Cluster C displayed a more balanced sex distribution (females n = 121, 52%) and was characterized by elevated frequencies of CD8+ T cells and NK cells, see **Figure 4b, c** and **Supplementary Figure 3a**. These differences were also evident when examining the immune cell modules and their associated gene expressions (**Figure 4d, e**). Specifically, Cluster A showed up-regulation of B cell and CD4+ T cell modules at both cellular and gene expression levels. Cluster B exhibited elevated levels in classical monocyte/mDC and nonclassical/ intermediate monocyte modules. Cluster C showed higher levels in the cytotoxic and CD8+ T cell modules, **Figure 4d, e**. Differentially gene expression analysis was employed to investigate the molecular differences across the three clusters. In total, 584 genes were up-regulated in cluster A, 1,411 genes in cluster B, and 786 genes in cluster C. Pathway enrichment analysis revealed that up-regulated genes in Cluster A and Cluster C were linked to activation of adaptive immune responses, whereas genes up-regulated in Cluster B were associated with inflammatory responses, **Supplementary Figure 3b-g**.

**Figure 4:**
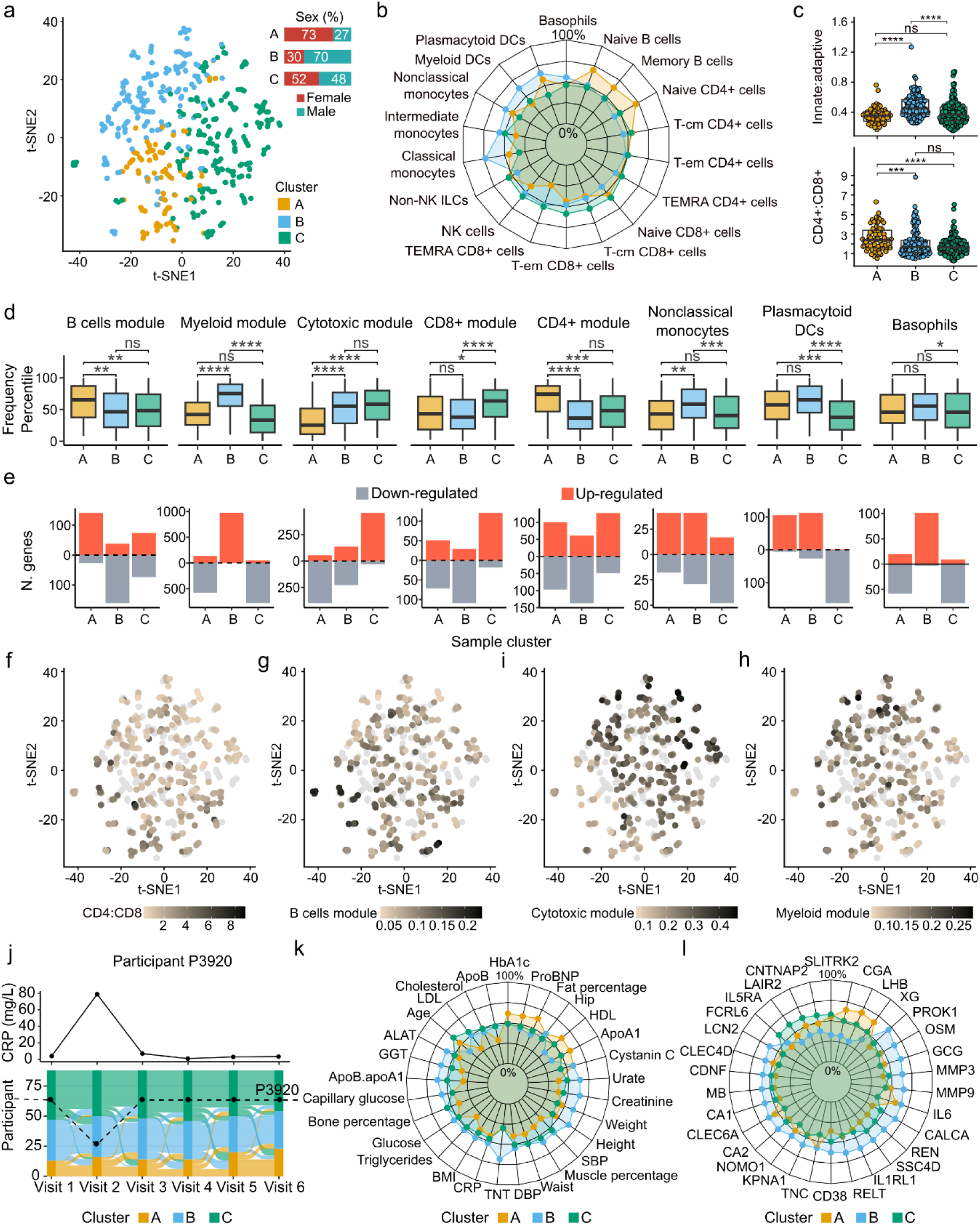
Sample clustering. (a) t-SNE sample clustering based on transcriptomic profiles of genes in the PBMC association network. (b) Immune frequencies patterns across the three sample clusters. (c) Distribution of (top) innate immune cell frequencies and (bottom) CD4:CD8 ratio across the three clusters. (d) Immune frequencies of the modules in the PBMC network, divided by sample clusters. (e) Number of up- and down-regulated genes in each module (FDR<0.05) obtained from differential expression analysis between each cluster and the remaining two. (f-h) t-SNE clustering colored based on CD4:CD8+ T cells ratio, and immune frequencies of the B cells, cytotoxic and myeloid modules. (j) (Top) CRP levels of participant P3920. (Bottom) Sample clusters of participants across visits; highlighted in black are the samples from participant P3920. (k) Pattern of clinical variables across the three clusters. (l) Patterns of the 30 proteins with the most significant up-regulation in any of the three clusters. SBP, systolic blood pressure; DBP, diastolic blood pressure; TNT, troponin T; CRP, C-reactive protein; HDL, high density lipoprotein; ALAT, alanine aminotransferase; GGT, gamma-glutamyl transferase. P-values are calculated by Wilcoxon tests in c-d; ns, not significant; * *P*< 0.05; ** *P* < 0.01; *** *P* < 0.001; **** *P* < 0.0001.

We next examined the longitudinal stability of the three immunotypes in samples across individuals in the S3WP cohort. Interestingly, we observed that samples collected from the same participants were mostly assigned to the same cluster through the six visits, reflecting the stability of individual’s immunotype, **Figure 4j**. In fact, around 65% of the participants had at least five of six samples were assigned to the same cluster. A notable exception was participant P3920, who experienced systemic inflammation at the second visit and accordingly shifted from cluster C to cluster B at that time point, before returning to cluster C at subsequent visits, **Figure 4j**.

Further characterization of the immunotypes using clinical variables and plasma proteomics revealed significant metabolic and inflammatory differences across the three clusters, **Figure 4k and 4l**, **Supplementary Table 4.** Overall, Cluster A exhibited a systemic profile typically associated with lower cardiometabolic and inflammatory risk, characterized by higher HDL and ApoA1, lower LDL, ApoB, triglycerides, urate, creatinine, CRP, as well as systolic and diastolic blood pressure. Cluster B, in contrast, showed clear pro-inflammatory signatures, including higher levels of CRP, IL6, GGT, TNT, and creatinine. Cluster C showed a relatively low inflammatory profile similar to Cluster A, but with higher metabolic risks. Compared to Cluster A, Cluster C demonstrated less favorable lipid parameters, including lower HDL and ApoA1 and higher LDL and ApoB (**Figure 4k**). Similar patterns were observed in the plasma proteome profiles, **Figure 4l, Supplementary Table 4**. Cluster A was characterized by elevated levels of several endocrine- and metabolism-associated proteins (LEP, LPL, GH1, LHB, CGA). The proteomic profile of Cluster B indicated a persistently elevated inflammatory milieu associated with higher cardiometabolic risk, characterized by increased levels of pro-inflammatory cytokines and chemokines (IL6, OSM, CXCL10, CXCL13, TREM1), acute phase markers (LCN2, SAA4, CES1/2), stress-related metabolic hormones (ANGPTL4, AGRP, FGF19, GCG), and markers of endothelial dysfunction (MMP3, MMP9, SERPINE1, VEGFC). On the other hand, Cluster C did not show enrichment for proteins related to inflammation or cardiometabolic health, but was instead characterized by immune-related programs, including lymphoid-associated receptors (FCRL5, FCRL6, CRTAM, CD48).

### Genetic architecture underlying inter-individual variation in immunotypes

To investigate whether genetic variation contributed to the individual differences in immunotypes, two separate genome-wide association studies (GWAS) were performed between immune cell frequencies and PBMC gene expression levels to 6.7 million common genetic variants (minor allele frequency, MAF > 0.05). In total, 319 significant associations between immune cell frequencies and genetic variants were identified (conventional *P* < 5 × 10^-8^ / 53). Among these, 39 independent (linkage disequilibrium, LD, r^2^ < 0.1, conditional *P* < 0.1) cell type quantitative trait loci (ctQTLs), significantly associated with 15 immune cell subtypes, were identified. The most evident association was linked to the frequency of memory CD4+ CD57-/CD39+ cells, a subset of Treg cells, and the gene *ENTPD1* (**Figure 5a**). *ENTPD1* encodes a plasma membrane protein that hydrolyzes extracellular ATP and ADP to AMP, and this association has been previously reported ^19,32,34^.

**Figure 5:**
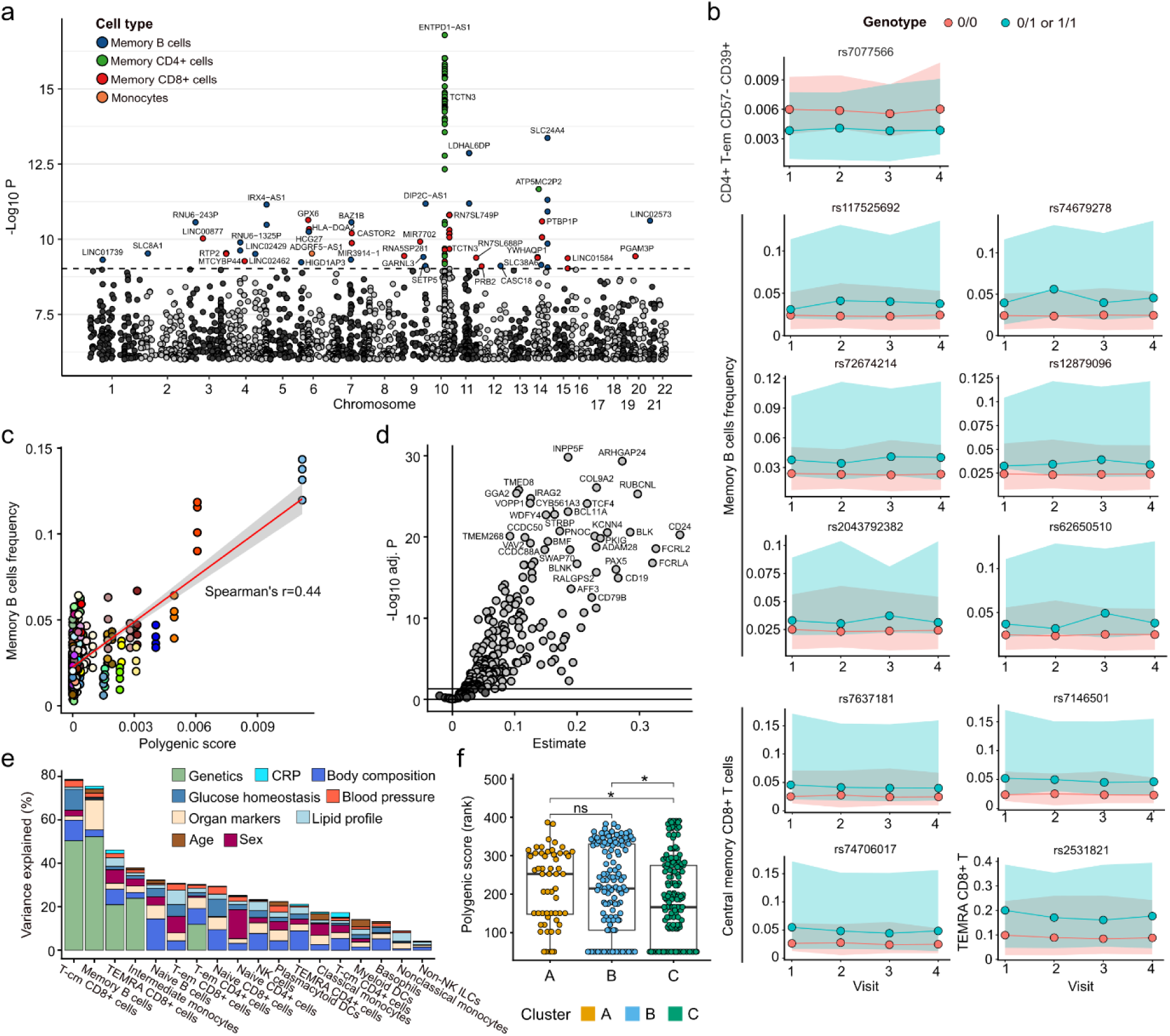
Analysis of genetic variation within the cohort. (a) Genome wide association analysis of 53 immune frequencies, colored by the corresponding immune family. (b) Examples of genetic variants association with increased memory CD4+ T cells, memory B cells and memory CD8+ T cell frequency. (c) Correlation between genetic score and frequency of memory B cells. (d) Volcano plot computed from the association between the polygenic score and the expression of genes associated with memory B cells in the PBMC association network. (e) Contribution of genetic and non-genetic factors to immune cell frequencies. (f) Polygenic score of memory B cells across the three sample clusters. Wilcoxon test was used for statistical analysis; ns, not significant; * *P* < 0.05.

Most of the genetic associations with immune cell frequencies (38 out of 39) were linked to adaptive immunity cell subtypes, primarily memory B cells (n = 19, 49%) and memory T-cm CD8+ cells (n = 14, 36%) (**Figure 5b** and **Supplementary Table 5**), while no ctQTLs were connected to naïve cells. The median value of the minor allele frequency (MAF) of ctQTLs associated with memory B cells was 0.054 (range 0.051–0.074), while the median for central memory CD8+ T cells was 0.064 (range 0.054–0.45). For example, the frequency of switched memory B cells with phenotype CD27+/CD22-/CD24-was linked to the highest number of genetic variants (n = 12, 31%), including <u>rs12879096</u> (intron variant of *SLC24A4*), <u>rs1228714111</u> (intron variant of *DIP2C*), and <u>rs1225770470</u> (intron variant of *BAZ1B*). GWAS on gene expressions confirmed genetic regulation of memory B cells; in fact, the genes associated with at least one independent eQTL were found to be enriched for genes that were significantly associated with memory B cells from the earlier analysis (Fisher’s exact test, p-value = 0.01, odds ratio = 1.8).

To investigate whether the identified genetic variants could explain the differences in immune frequencies in the cohort, we computed the polygenic score (PGS) of memory B cells. We observed significant correlation (Spearman’s ρ = 0.44) between PGS and frequency of memory B cells, **Figure 5c**. Moreover, we also observed a significant correlation between PGS and the gene expression of genes in the B cell module, **Figure 5d**. Notably, 243 out of 249 genes showed a positive association, out of which 209 had a significant p-value (FDR = 0.05), underscoring the genetic regulation on both B cell frequencies and gene expression. To compare the influence of genetic and non-genetic variables on immune frequencies, we performed ANOVA, which highlighted T-cm CD8+ T cells and memory B cells as the immune populations most regulated by genetics, **Figure 5e**. On the other hand, clinical variables had a more even contribution across cell types, explaining an average of around 21% of the variance in immune cell frequencies, with the largest contributions arising from body composition and organ-related biomarkers. Lastly, we also observed statistically significant differences in PGS across the three sample clusters, namely Cluster C was characterized by significantly lower PGS compared to the remaining two, **Figure 5f**.

## Discussion

In this study we provided a longitudinal, systems-level characterization of immune variation across 101 healthy individuals. By integrating four paired omics (PBMC transcriptomics and mass cytometry, whole genome sequencing and plasma proteomics), we were able to characterize individual immune features across multiple layers and leveraged cross-omics validation to increase confidence in the biological validity of our results. Using repeated measurements from the same individuals, we investigated the presence of stable molecular signatures across time, characterized by high between- and low within-subject variability ^35,36^, an approach that has been used to identify immune markers that characterize the predisposition of the individual to differential immune responses ^37^. Among the genes with highest inter-individual differences, we could identify genes that were previously shown to drive individual differences in immune responses. For example, polymorphisms in the HLA-I and HLA-II loci are hallmarks of autoimmune disease risk ^38,39^; *ERAP2* and *SIGLEC14* have been associated with genetic variants (*ERAP2*: rs2549794, rs2248374; *SIGLEC14*: null/fusion allele) that predict respiratory infection severity, and are thus linked to individual immune responses ^40,41^; genetic variations in *GPSM3* (rs204989 and rs204991) have been linked to differential gene expression and increased risk of rheumatoid arthritis ^42^.

The individuality of the human immune system was explored across four levels of resolution. First, using dimensionality reduction techniques, broad molecular patterns that distinguish individual immune profiles. The strong individual clustering observed in transcriptomics data indicated that each participant possessed a unique immunotype. Remarkably, even after one year, every sample could be unequivocally mapped to the correct participant from measurements taken the previous year. Secondly, we assessed inter-individual variability across gene expression, immune cell frequencies, and protein abundance, revealing consistent individual-specific patterns across all three layers that converged on shared immune functions. Genes with high individual variability included those involved in antigen processing and presentation (HLA genes, *ERAP2*, *TAPBP*, *PSMB9*), cytotoxic effector functions (*CD8A*, *GZMH*, *GNLY*), inflammasome modulation (*SIGLEC14*, *TMEM176A/B*), and B-cell regulation (*FCER2*, *GPSM3*, *ZNF683*). Proteins mirrored these pathways, highlighting innate sensing and myeloid activation (TLR3, CLEC7A, CHIT1, FOLR3, MBL2), stress-induced cytotoxic ligands (MICA/B), immune regulatory pathways (LAIR2, LILRB5, FCRL family, CD6, CD70, PILRB), and additional B-cell–associated molecules (FCRLB, FCRL1, FCRL6, CD70). At the cellular level, memory T- and B-cell subsets, as well as NK cells, exhibited the highest inter-individual variability in frequency, reflecting the compartments through which these molecular programs are enacted. Thirdly, to conduct a systematic analysis on inter-individual differences centered around immune populations, we performed a genome-wide association analysis between transcripts and immune frequencies. A thorough analysis highlighted three components of the network characterized by shared biological functions and strong individuality: B cells, cytotoxic cells (NK and TEMRA CD8+ T cells) and myeloid cells (classical monocytes and myeloid DCs). Fourthly, GWAS on immune cell composition and gene expression were performed to investigate differences in genotype across the cohort. Independent GWAS of the two omics layers revealed a strong genetic regulation of memory B cells, affecting both their frequency and transcriptional profiles. Moreover, a polygenic score for memory B cell frequency showed consistent genetic effects on both cell abundance and gene expression, reinforcing the presence of robust genetic control.

Our analyses suggested that individual differences were centered around three major immune cell types: B cells, cytotoxic cells (NK and TEMRA CD8+ T cells) and myeloid cells (classical monocytes and myeloid DCs). Accordingly, sample clustering of the transcriptomic profiles confirmed that these three components captured the primary sources of variation within the cohort, defining three distinct immunotypes with well-characterized molecular and clinical features. Cluster A was associated with higher frequency and gene expression of B cells, which were largely controlled by genetic predisposition. The samples within this cluster consisted predominantly of females with favorable metabolic indicators. On the other hand, samples in Cluster B were composed mainly of males and were characterized by higher activation of classical myeloid functions. In accordance with the functions of these cell types, these samples showed higher inflammation markers and pro-inflammatory cytokines in the circulating proteins, indicative of a state of chronic inflammation. Lastly, Cluster C showed less favorable metabolic markers than Cluster A, coupled with increased cytotoxic activity and cytokine signaling in circulating plasma. Overall, these results are in line with a recent study by Smithmyer et al. ^43^, which identified that CD8+ T cell activity and innate immune activation as two major axes of variation within a cohort of healthy individuals. Taken together, our analyses suggest that healthy immunotypes can be stratified into biologically meaningful groups, each reflecting an interplay between sex, genetic variants and metabolic health. Importantly, identifying such distinct immune profiles has direct implications for precision medicine, as it highlights the main axes of immune variation between individuals, providing a framework to understand individual immune responses. Although genetic effects on immune traits were detectable and consistent across immune cell composition and gene expression, only modest differences in genotype across the three clusters were identified, suggesting that the observed immunotypes were shaped primarily by environmental and cellular factors, rather than being strongly driven by genetic variation.

However, several limitations in the current study should be acknowledged. First, GWAS analyses were limited by the relatively small cohort, restricted to middle-aged Scandinavian adults, limiting statistical power and generalizability to other population groups. Secondly, the participants were relatively close in age (50-65 years old at the first visit), which likely led to an underestimation of the effects of age, a potent regulator of the immune system ^44^. Thirdly, the study included only healthy participants, preventing analysis of disease incidence or severity in relation to individual immunotypes. The observations made in this study remain descriptive and do not establish causal relationships or functional consequences. Further longitudinal and functional studies will be required to determine whether these stable, person-specific immune features contribute to disease susceptibility.

In conclusion, our study provided a longitudinal, multi-layered view of human immune variability, showing that individual immune profiles were relatively stable over time and organized into coordinated cellular and molecular modules. These modules delineated major immunotype axes that were associated with systemic physiological states and showed evidence of partial genetic influence. Together, these findings contributed to a systems-level understanding of baseline immune heterogeneity in humans and provided a basis for future studies in precision immunology.

## Materials and Methods

### Experimental Design

The objective of this study is to investigate immune system individuality in healthy individuals by integration of multi omics data. The data was retrieved from The Swedish SciLifeLab SCAPIS Wellness Profiling (S3WP) program, a non-interventional initiative aimed at gathering longitudinal clinical and molecular data from a community-based cohort. This program builds upon the Swedish CArdioPulmonary bioImage Study (SCAPIS), which is a prospective observational study encompassing 30,154 individuals aged 50 to 65 years at enrollment, randomly selected from the general Swedish population between 2015 and 2018 ^45^. The S3WP study consists of 101 healthy individuals who were previously extensively assessed in SCAPIS. Throughout the duration of the study, follow-up visits are conducted every third month (± 2 weeks) in the first year and approximately a 6-month interval in the second year. At each follow-up, blood samples were collected from individuals, along with anthropometric measurements, clinical chemistry, and hematological measurements, among other indicators. Whole genome sequencing data were detected at the baseline. Proteomic data were examined at each follow-up visit.

The study has been approved by the Ethical Review Board of Göteborg, Sweden (registration number 407-15), and all participants provided written informed consent. The study protocol adheres to the ethical guidelines outlined in the 1975 Declaration of Helsinki.

### Mass-cytometry data

Cryopreserved PBMCs were thawed in RPMI medium supplemented with 10% FBS, 1% penicillin streptomycin, and benzonase (Sigma-Aldrich). Cells were resuspended in RPMI medium supplemented with 10% FBS and 1% penicillin streptomycin and rested overnight at 37C in 5% CO2 for cells to be revitalized. Samples were randomized in batches to avoid systematic technical variation due to PBMC sample processing ^46^. Overnight rested cells were counted and checked for their viabilities. For live-dead discrimination, cells were stained with 2.5 mM Cisplatin (Fluidigm Inc.) in RPMI without FBS for 5 min at room temperature, followed by quenching with RPMI containing FBS. Cells were then fixed with 1% formaldehyde (Polysciences Inc.), washed, and resuspended in Cell staining buffer (CSB) (PBS with 0.1% BSA, 0.05% sodium azide and 2mM EDTA). For surface marker staining, cells were incubated for 30 min at 4C with a 30ul cocktail of metal conjugated antibodies (Antibodies listed in Key Resources Table) targeting the surface antigens, washed with CSB and fixed with 4% paraformaldehyde, all of which performed using a custom-built liquid handling robotic platform ^47^. Cells fixed in 4% paraformaldehyde were stained with Iridium-labeled DNA intercalator at a final concentration of 0.125 mM (MaxParâ Intercalator-Ir, Fluidigm Inc.) on the day of sample acquisition. Following incubation for 20 min at room temperature, cells were washed with CSB twice, once with PBS, and twice with milliQ water. Cells were counted and diluted to 500,000 cells/ ml in milliQ water containing 10% EQ Four Element Calibration Beads (Standard BioTools, formerly Fluidigm Inc.) and filtered through a 35mm nylon mesh ^48^. Samples were acquired on one of two CyTOF2 mass cytometers (Standard BioTools) using CyTOF software version 6.0.626 with noise reduction, a lower convolution threshold of 200, event length limits of 10-150 pushes, a sigma value of 3, and flow rate of 0.045 ml/min. All FCS-files unrandomized using the CyTOF software (version 6.0.626) were transferred without any additional preprocessing. Files were normalized using our own in-house implementation of normalization software described previously ^49^. Cells populations were manually gated using a Brodin lab developed software, Cytopy. Cell frequencies were batch corrected using the ComBat algorithm provided in the sva R package. To ensure the data was compositional, negative values were replaced with 2.2 × 10^-16^ and the sum of counts of each sample was normalized to the value of 1.

### RNA sequencing data

Total RNA was extracted using RNeasy Mini Kit (Qiagen) and quantified using Qubit 2.0 Fluorometer (Invitrogen). RNA was converted to a sequencing library using the TruSeq Stranded mRNA HT library preparation method (Illumina) using 500 ng of total RNA as input quantity. The obtained library was quantified using Qubit Broad Range assay kit or Quant-IT RiboGreen chemistry (Invitrogen). The obtained libraries were sequenced using Hiseq 2500 (Illumina) using either pair-end 100 bp or pair-end 125 bp in rapid run mode or high output mode, respectively. Each sample was sequenced targeting 30 M read pairs. Demultiplexing was done without allowing any mismatches in the index sequences. To obtain quantification scores for all human genes and transcripts across all samples, transcript expression levels were calculated as transcript per million (TPM) by mapping processed reads to the human reference genome GRCh37/hg19 and with gene models based on Ensembl (v92) ^50^ using Kallisto (v.0.43.1) ^51^. Data from multiple visits were integrated using batch correction implemented as removeBatchEffect in the R package limma v. 3.64.3 ^52^ using the sampling date as a batch parameter. This resulted in gene expression data for 19,670 genes out of which 8,368 were retained after filtering using the function filterByExpr from the R package edgeR v. 3.40.2 (with arguments min.count = 5, min.total.count = 10).

### Plasma protein profiling

We used a multiplex proximity extension assay (Olink Bioscience, Uppsala Sweden) to measure the relative concentrations of 794 plasma proteins in eleven Olink panels. To minimize inter-run and intra-run variation, the samples were randomized across plates and normalized using both an internal control (extension control) and an inter-plate control; then a pre-determined correction factor was applied to transform the data. The pre-processed data were provided in the arbitrary unit Normalized Protein eXpression (NPX) on a log2 scale. QC procedures were performed at both sample and protein level. Briefly, samples were flagged (did not pass QC) if the incubation control deviated more than a pre-determined value (+/− 0.3) from the median value of all samples on the plate (www.olink.com). To reduce the batch effect between samples run at different times, bridging reference samples from different visits were also run on plates from the different batches. Reference sample normalization based on bridging samples was conducted to minimize technical variation between batches (www.olink.com). After QC, a total of 794 unique proteins for 90 subjects and 6 visits (540 samples) were retained for analysis. The detailed information about plasma protein profiling can be found in previous papers ^53,54^.

### PBMC network construction

To identify significant associations between gene expression and immune frequencies, we performed a systematic regression analysis across all measured transcripts and immune frequencies. The underlying hypothesis is that the measured expression of each transcript was given by the sum of its expression across cell types. Given the compositional nature of immune frequencies, specific methods must be used to obtain robust results. In this work, we have adopted the strategy proposed by Hron et al. ^30^ and adapted it to include repeated measurements. To estimate the expression of each measured gene across PBMC immune populations, we built a linear model where gene expression was the dependent variable and all immune cell frequencies the explanatory variables. Given the compositional nature of the explanatory variables, *i.e.*, immune cell frequencies, we adopted a method based on recursive isometric logratio (ILR) transformations ^30^. To assess the relationship between a given gene and PBMC frequencies, represented by the vector

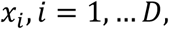

where D is the number of immune populations, we used the following ILR transformation:

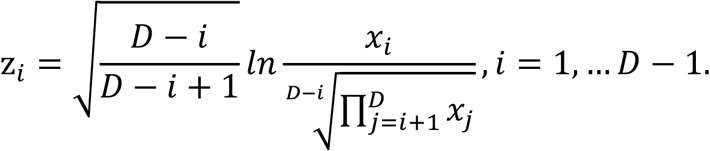

After performing standard linear regression between the gene expression and the vector z, its first component z_1_, will be interpretable as the association between the first immune cell frequency x_1_ and gene expression. Therefore, for every gene, the procedure was repeated by shifting the positions of vector x, to obtain interpretable estimates for all populations. Sex and age were included as covariates in the model. This regression analysis was performed independently for each visit, the resulting estimates were then combined using a random-effects meta-analysis to account for between-visit variability, using the function rma from the package metafor v. 4.8.0.

### Validation of the gene-cell type associations

The associations in the network were validated using multiple approaches. The Human Protein Atlas v.23 was accessed to retrieve known associations between genes and immune cell types (“elevated” genes) based on 18 immune cell populations isolated by flow cytometric sorting (https://www.proteinatlas.org/humanproteome/single+cell/immune+cell). Fisher’s exact tests were performed between each module in the network and population in the annotation to identify significant overlaps; p-values were corrected using the Benjamini-Hochberg procedure. Secondly, we merged two lists of PBMCs markers: Azimuth ^31^ (https://azimuth.hubmapconsortium.org/references/) and DICE (Database of Immune Cell Expression, eQTLs, and Epigenomics) ^32^, available at https://dice-database.org. Moreover, we downloaded the transcriptomic profiles available from the Human Protein Atlas (https://www.proteinatlas.org/humanproteome/blood) and used them to verify the expression patterns of key genes across cell types.

Gene ontology enrichment analysis and gene set enrichment analysis were performed using the R functions enrichGO and gseGO from the package clusterProfiler v. 4.6.2.

### Sample clustering algorithm

The K Nearest Neighbour Search (KNN) algorithm was implemented to calculate the adjacency matrix, using the nn2 function from the R package RANN v2.6.1 ^55^, with the maximum number of nearest neighbors to compute set to 15. An adjacency matrix was built based on the result of the nearest neighbors of each metabolite and used to create a graph using the R package igraph v1.5.0 ^56^. The Louvain algorithm was used to find communities in the graph, implemented by the function cluster_louvain from the R package igraph ^56^. The alluvial plot was created using the R package ggsankey v. 0.0.99999. Radar charts were created using the R package fmsb v. 0.7.6.

### Genome-wide association analysis

The whole genome-sequencing procedure has been detailed previously in Zhong et al. ^53,54^. Briefly, Genomic DNA was sequenced to average 30X coverage on the HiSeq X system (Illumina, paired-end 2 × 150 bp). The alignment was performed using BWAmem, reference genome GRCh38.p7. Single-nucleotide and insertion/deletion variants were called following the GATK pipeline (https://software.broadinstitute.org/gatk/best-practices; GATK v3.6). BCFtools ^57^ and plink 2.0 ^58^ were used to perform quality control. The exclusion criteria included: (1) removing variants which did not receive the “PASS” tag from GATK; (2) removing variants with minQUAL < 30; (3) removing variants/samples that with a genotyping rate < 0.05; (5) removing variants with a low minor allele frequency (MAF) (< 5%); (6) removing variants that failed the Hardy–Weinberg equilibrium (HWE) test (*P* < 1×10^-6^). In total, 6,691,390 high-quality variants were identified in all samples.

GWAS were performed to identify significant genetic associations to immune cell frequencies and gene expression; we used linear models adjusted for sex and age (at baseline) to fit to the mean values of immune cell frequencies and gene expression across visits. All the GWAS analysis were performed using PLINK v2.0 ^58^. P*-*values were corrected for multiple hypothesis testing using the Bonferroni correction (*i.e.*, 5 × 10^-8^ / 53 for immune frequencies and 5 × 10^-8^ / 8,368 for gene expression).

To identify independent ctQTLs/eQTLs for a given metabolite, linkage disequilibrium (LD) *r*^2^ > 0.05 with window size 500 kb was first used to exclude the correlated variants. When multiple ctQTLs/eQTLs were associated to the same frequency/gene, a conditional analysis was then carried out in which the genetic associations were re-calculated using the lead SNP as covariate. Only associations with conditional P-value < 0.1 were considered to be independent ctQTLs/eQTLs. SNPs were annotated to the closest gene using the function getBM from the R package biomaRt v. 2.64.0.

## Statistical analysis

T-distributed stochastic neighbor embedding (t-SNE) was performed to visualize similarities between samples, using the R package Rtsne v.0.17. To equalize the number of variables across different omics, PCA was performed on each dataset, retaining the first 50 principal components that were subsequently used as input to generate t-SNE.

We parsed CyTOF data and extracted measurements from 10 cells per sample and immune population, which were further filtered by selecting 1,000 random measurements per immune population. The data were transformed using an inverse hyperbolic sine function, scaled by a factor of 1/5, and standardized (Z-transformed). Uniform manifold approximation and projection was performed using the R package umap v. 0.2.10.0. Hierarchical clustering was performed using the average cell surface protein expression of each cell type, implemented by the function ‘hclust’ (agglomeration method ‘complete’).

Differential analyses were performed to identify statistically significant differences in gene expression and protein abundance between the three clusters. Differential analysis was performed using the package limma v. 3.64.3. Contrasts were defined to compare each cluster to the mean gene expression/protein abundance of the other two clusters. P-values were adjusted using the Benjamini-Hochberg procedure and a threshold of FDR = 0.05 was used to classify statistically significant results. Similarly, to investigate differences in clinical variables, we performed univariate linear modeling. Contrasts were implemented using the function ‘emmeans’ from the package emmans v. 2.0.0 (method ‘del.eff’).

To calculate the polygenic score (PGS) of memory B cells frequency, the alternative allele dosage of every B cell subtype of each individual was multiplied by the estimated coefficient from GWAS. The value was adjusted by the relative frequency of the cell subtype. Linear regression was used to estimate the relationship between the memory B cell PGS and the expression of genes included in the B cell module of the PBMC network.

## Acknowledgments

The processing of whole genome sequencing data was performed on resources provided by SNIC through Uppsala Multidisciplinary Center for Advanced Computational Science (UPPMAX) under Project SNIC 2017/7-342 and the integration of multi-omics was performed under Project sens2018115.

## Author contributions

WZ conceived and designed the study. MU, GB, FE, TL, JM, and LF, collected and contributed data to the study. AZ, XW, ZT, YC, and WZ performed the data analysis. GB supplied clinical material. AZ, XW and WZ drafted the manuscript. All authors read and approved of the final manuscript.

## Data and materials availability

This study utilizes participant-level datasets that have been securely deposited at the Swedish National Data Service (SND), a repository certified by the Core Trust Seal (https://snd.gu.se/). In compliance with patient consent and confidentiality agreements, these datasets are accessible for validation purposes only. Requests for access can be directed to SND via email. The evaluation of such requests will be conducted in accordance with relevant Swedish legislation. Additionally, for inquiries specifically on research within the scope of the S3WP program, interested parties are encouraged to contact the corresponding author directly. The code is available at the GitHub page https://github.com/Wen-Zhong-Lab/Individualized-immunotypes.

