## Supplementary Figures for "Systems-level longitudinal immune profiling reveals individualized immunotypes and genetic associations"

Alberto Zenere *et al.*

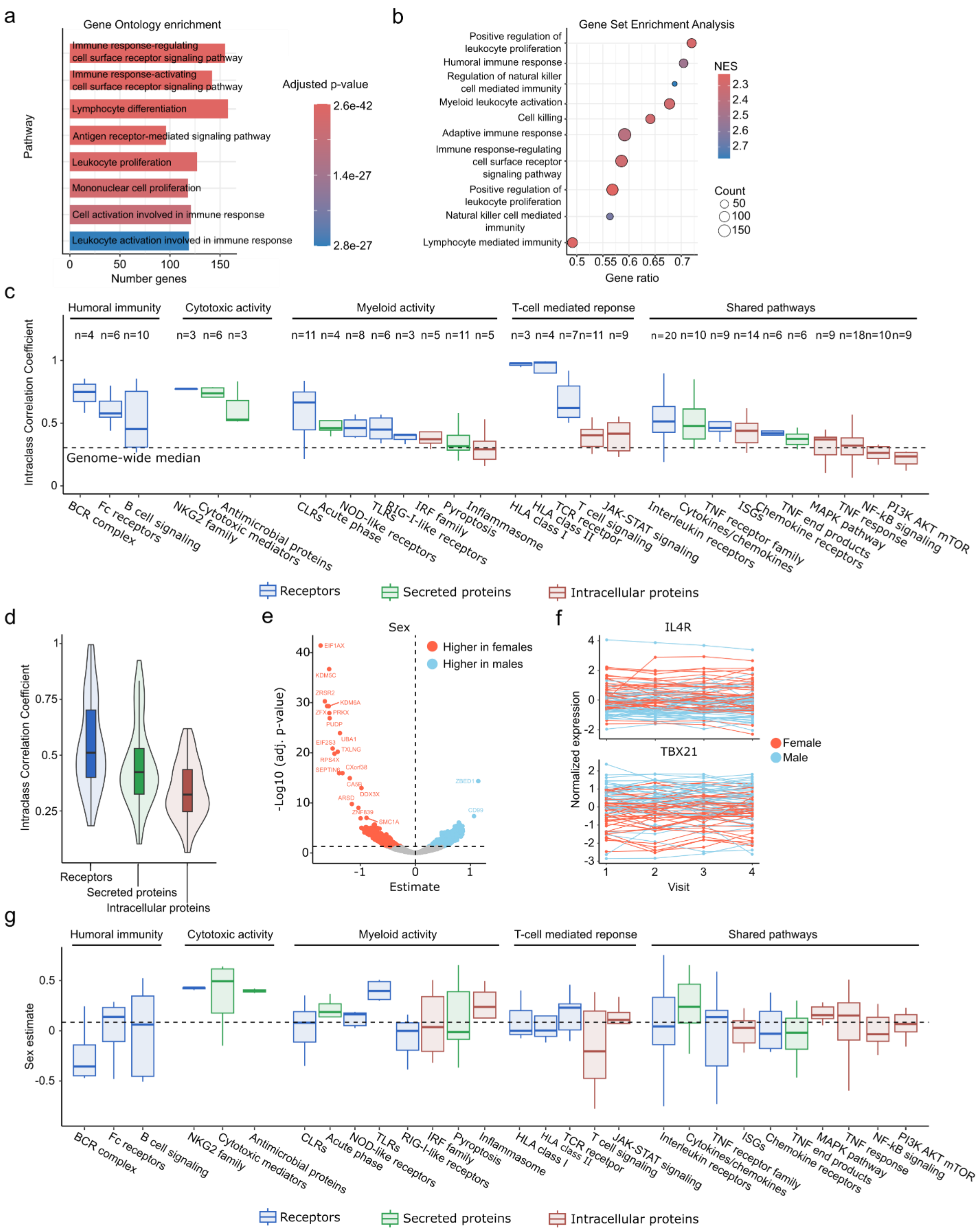


**Supplementary Figure 1.** (a) Pathway enrichment analysis of genes with higher inter- than intra-individual variation. (b) Gene set enrichment analysis of all 8,368 measured genes ordered based on their intraclass correlation coefficient. (c) Intraclass correlation coefficient of 213 genes that were manually categorized into immune pathway. (d) Distribution of gene intraclass correlation coefficients, based on the protein function: receptor, secreted protein or intracellular protein. (e) Volcano plot representing sex effect on gene expression, using the p-values and coefficients obtained from the calculation of intraclass correlation coefficients. (f) Examples of genes with significant sex and individual effect. Red denotes samples from females, while light blue from males. (g) Sex effect of genes grouped by immune pathway.

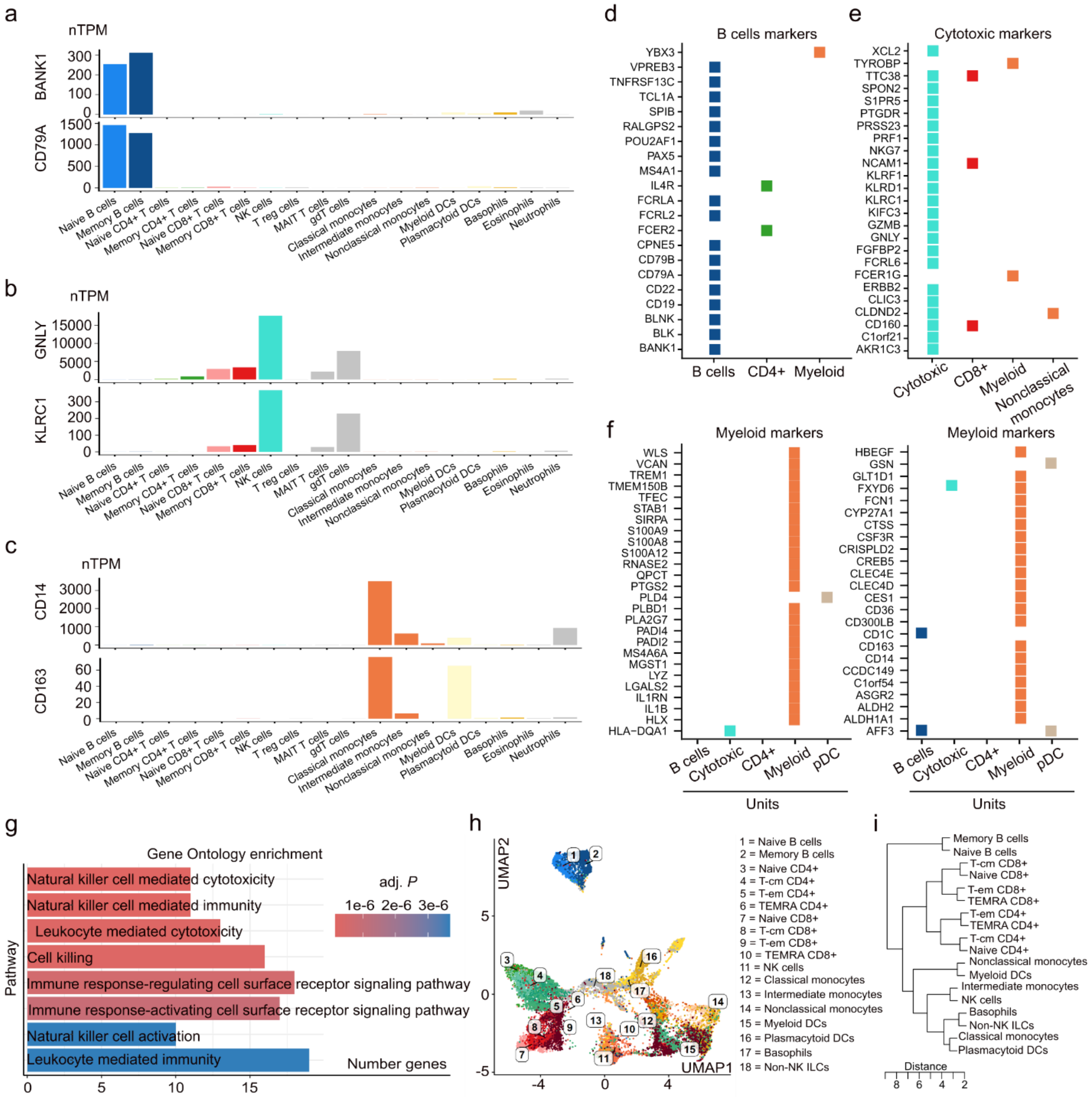


**Supplementary Figure 2.** (a-c) Examples of expression profiles of two B cell markers, cytotoxic cell markers and myeloid cell markers, using transcriptomic data from the Human Protein Atlas. (d-f) Associations within the PBMC network of known markers retrieved from Azimuth and DICE. (g) Gene Ontology enrichment of genes significantly associated with NK cells and TEMRA CD8+ T cells. (h) Uniform manifold approximation and projection based on single cell protein surface markers measured by CyTOF. (i) Hierarchical clustering of PBMC immune populations based on surface protein markers measured by CyTOF.

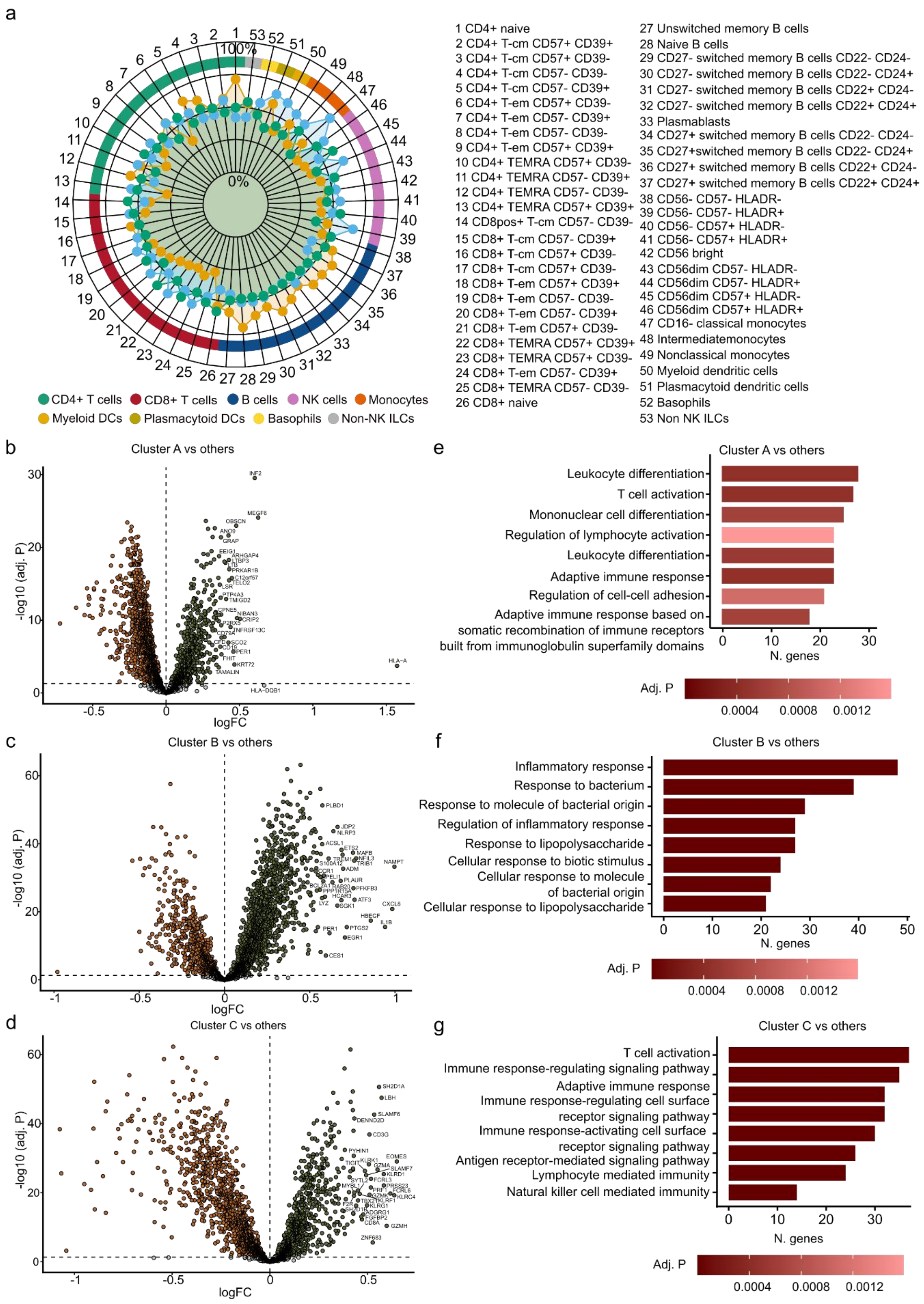


**Supplementary Figure 3.** (a) Immune frequency patterns of the 53 measured cell types. (b-d) Gene expression differential analysis. (e-g) Pathway enrichment analysis of the significant (FDR=0.05) genes.
